# SURFMAP: a software for mapping in two dimensions protein surface features

**DOI:** 10.1101/2021.10.15.464543

**Authors:** Hugo Schweke, Marie-Hélène Mucchielli, Nicolas Chevrollier, Simon Gosset, Anne Lopes

**Author notes:** **Corresponding Author** Anne Lopes, Université Paris-Saclay, CEA, CNRS, Institute for Integrative Biology of the Cell (I2BC), 91198, Gif-sur-Yvette, France., Hugo Schweke, Department of Chemical and Structural Biology, Weizmann Institute of Science, Rehovot 7610001, Israel.

## Abstract

Molecular cartography using two-dimensional (2D) representation of protein surfaces has been shown to be very promising for protein surface analysis. Here, we present SURFMAP, a free standalone and easy-to-use software that enables the fast and automated 2D projection of either predefined features of protein surface (i.e., electrostatic potential, Kyte-Doolittle hydrophobicity, stickiness, and surface relief) or any descriptor encoded in the temperature factor column of a PDB file. SURFMAP uses a pseudo-cylindrical sinusoidal “equal-area” projection that has the advantage of preserving the area measures. It provides the user with (i) 2D maps that enable the easy and visual analysis of protein surface features of interest and (ii) maps in a text file format allowing the fast and straightforward quantitative comparison of 2D maps of homologous proteins.

## INTRODUCTION

Genome sequencing has produced a huge amount of protein sequences and motivated the development of methods for protein comparison and classification. In particular, important efforts have been made to predict the function of a protein of interest from the comparison of its sequence with those of already annotated homologs. If most methods focus on the information extracted from protein sequences, large-scale structural genomic experiments have promoted the development of three-dimensional (3D) structure comparison-based methods, which are very powerful when comparing remote homologs^1–4^. Indeed, protein structure is more conserved than protein sequence and these methods are able to highlight proteins displaying similar structure while sharing little sequence identity. Nevertheless, most sequence-based and structure-based methods extract information from the entire protein. It is thus difficult to distinguish residues conserved to maintain protein function from those ensuring protein stability. This is even more problematic when dealing with large families of paralogs that share the same fold but display different functions and/or physico-chemical properties. Precisely, being able to predict interaction preferences (protein partners, ligands, co-factors...), oligomeric states or to identify thermostable proteins, for instance, would be of great value for the classification and annotation of these large protein families.

Protein surfaces can be considered as a proxy for protein interactions, thereby playing a critical role in protein function^5^. Several methods have been developed to analyze and compare protein surface features. Some of them focus on specific regions of the surface such as protein binding sites, active sites, binding pockets^6–10^, and ignore the rest of the surface which plays an important role in protein interactions by constantly competing with the interaction sites^11^. On the other hand, “molecular cartography” which has been introduced by Fanning et al^12^ is based on a system dimension reduction and enables the representation of the whole protein surface in a synthetic and robust way. The principle consists in projecting the 3D structure of a protein into two dimensions (2D) and then mapping features of interest (charges, hydrophobic patches, topography, evolutionary conserved sequence conservation…) on the resulting 2D map. Despite the efforts made on protein 3D structure representation tools^13–15^, handling 3D objects is always difficult, and these 2D maps greatly ease the visualization and analysis of the distribution of a given feature. For example, they enable the visual comparison of the distribution on homologous proteins’ surfaces of specific features to understand the molecular basis of their common or different interaction properties. Additionally, 2D maps are well suited for large-scale protein surface comparisons^11^ through the calculation of a map similarity with a straightforward numerical measure, and since the dimension reduction is robust against local irregularities of protein surfaces.

Although “molecular cartography” is not recent and has been shown to be very promising for protein function annotation, only a few methods have been proposed so far. Godzik et al^5^ proposed a protein surface representation based on a spherical approximation of the protein surface in order to compare the distribution of physico-chemical features on the surface of homologous proteins. Other methods with interesting outcomes in visualization and/or surface feature comparison^16–20^ are restricted to a limited subset of surface descriptors or do not provide the tool to compute the corresponding 2D maps. Recently, Kontopoulos et al, developed Structuprint^21^, a program that enables the visualization of more than 300 protein surface features. They use the Miller cylindrical projection^22^, i.e. a projection that conserves the angles and the shapes of small surfaces but that induces an important overestimation of the area at the poles. This can be problematic since the different protein surface regions do not contribute equally when comparing different surface maps, and the assignment of surface regions either to the poles or equator results from the arbitrary orientation of the protein before projection.

Here, we present SURFMAP, a program that enables the projection of predefined protein surface features: (i.e. electrostatic potential, hydrophobicity, stickiness, surface topology), residues of interest, or any descriptor encoded in the temperature factor column of a PDB file. SURFMAP uses a pseudo-cylindrical sinusoidal equal-area projection that has the advantage of preserving the area measures at the cost of distorting shapes locally. Here, we aim to preserve the size of the regions of interest rather than providing a precise representation of their shapes. Indeed, we need for a method robust against local variations since (i) proteins are flexible objects and the shapes of surface features can vary locally, (ii) protein surfaces display many local irregularities that could introduce noise when comparing two homologous surfaces, and (iii), nowadays, we frequently use 3D models that can be more or less accurate depending on their similarity with the template used for building the model. We illustrate the application of SURFMAP with three different examples of analysis on the family of superoxide dismutases (SODs). These proteins are enzymes that catalyze the dismutation of the superoxide radical to hydrogen peroxide and dioxygen. We show how SURFMAP can be used to either compare the distribution of different surface features of a protein of interest or to compare the distribution of same surface features of different protein homologs. Interestingly, we show that the straightforward comparison of the 2D maps based on the stickiness feature (that reflects the interaction propensity of the protein surface) enables the distinction of SODs with different oligomerization states and different metal ions binding preferences.

## METHODS

### Calculation of 2D maps

The cartography of protein surface properties is done in 5 steps:

#### 1 Calculation of the protein surface features

The first step consists in calculating the values of the feature(s) of interest for each protein residue or atom (Figure 1A). When the feature is calculated at the residue scale, each atom receives the value of the residue corresponding residue. Five different features can be calculated by SURFMAP. The Kyte-Doolittle and Wimley White hydrophobicity are mapped at the residue scale and are directly derived from the Kyte-Doolittle^23^ and Wimley-White^24^ scales, respectively. The stickiness scale^25^ reflects the propensity of each amino acid to be involved in protein-protein interfaces. The circular variance^26^ (CV) can be calculated at the residue or atomic scale and measures the atom density around each atom. The CV provides a useful descriptor of the local geometry of a surface region. CV values range between 0 and 1 with low values reflecting protruding residues, and high values indicating residues located in cavities. The electrostatic potential of the protein surface is calculated at the atomic scale with the APBS software^27^. In addition, SURFMAP can handle any other feature stored in the temperature factor of the input PDB file.

**Figure 1:**
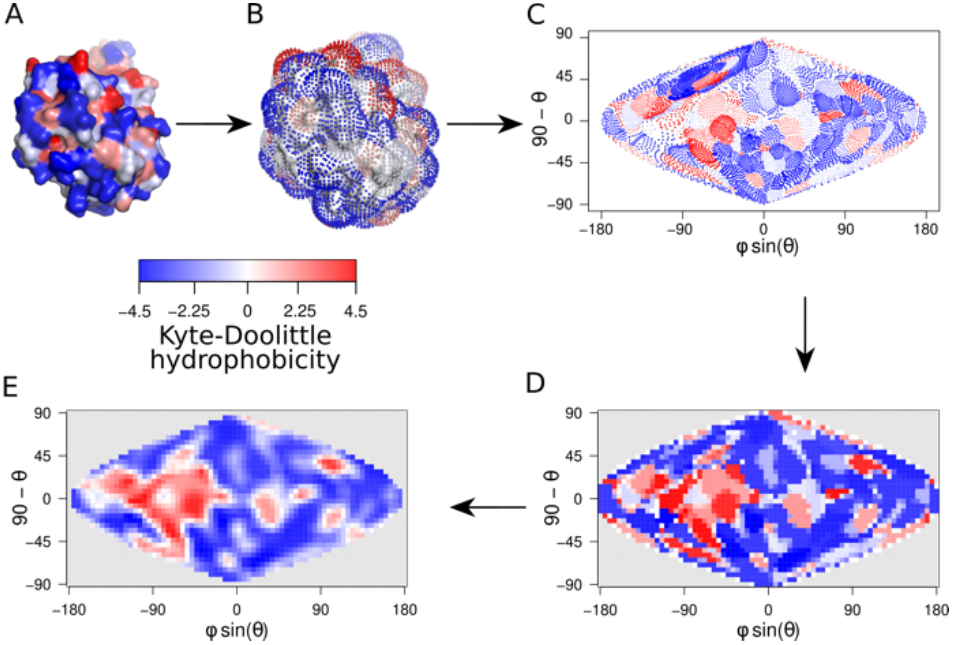
Generation of 2D maps. Generation of the Kyte-Doolittle hydrophobicity 2D map for the chain A of the cambialistic superoxidase dismutase of *P. Shermanii* (PDB code: 1ar4). (A) Each atom (or residue depending on the mapped property) is associated with its corresponding value and colored accordingly (KD hydrophobicity in this example). (B) Generation of a set of particles around the protein surface with MSMS^28^. Each particle is located at 3Å (default) from the closest residue of the protein of interest. The KD hydrophobicity value of the closest residue of the protein is then attributed to each particle. (C) The spherical coordinates (*φ*, *θ*) (with respect to the protein center of mass) are projected on a 2D map in an equal-area 2D sinusoidal projection. Each projected dot is then associated with the feature value of its corresponding particle. (D) The 2D map is divided into a grid of 36×72 cells and each cell is associated with the average of the corresponding values. (E) Cell values are then smoothed by averaging the value of each cell with the one of the eight surrounding cells. The same color scale is used for all panels (A-E).

#### 2 Generation of a set of points around the surface of the protein

The second step consists in generating a set of particles localized at 3Å (default value but this value can be modified by the user) from the surface of the protein of interest. This set of points is generated with MSMS^28^, a software that computes the molecular surface using a reduced surface representation of proteins (see^28^ for more details). Here we refer to this ensemble of points as the “shell” (Figure 1B). This shell enables the precise mapping of the protein surface on a 2D map. It provides more complete information than the direct mapping of the atoms, as in the latter case the number of grid cells usually largely outnumbers the number of atoms, which results in a lot of cells left unassigned and containing no information.

#### 3 Assignment of the value of the mapped feature to each particle of the shell

Each shell particle is annotated with the feature value of the closest protein atom (Figure 1B).

#### 4 Sinusoidal projection on a 2D plan of the spherical coordinates of each shell particle with respect to the center of mass of the protein

The coordinates of each shell particle *pi* are expressed in spherical coordinates (*φ*, *θ*) with respect to the center of mass of the protein noted *G. φ* is the angle between the Y axis and the projected vector of vector 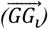 in the plan 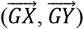, while θ is the angle between the vector 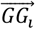 and the Z axis. The coordinates of the shell particles are then projected onto a 2-dimensional plane by a sinusoidal projection. Each particle is therefore represented on the 2-D plane by a couple of coordinates (x = ϕsin *θ*, y = 90 − *θ),* the latter being associated with the feature value of the corresponding particle (Figure 1C) (see Recio et al^29^ for more details).

#### 5 Division of the map and smoothing

The resulting map is then divided into 72 × 36 cells (the grid resolution can be modified by the user). The sinusoidal projection is equal-area. Consequently, each cell represents a surface of the same area. Each cell is then associated with the average of the particle values it contains (Figure 1D). The map can then be smoothed by averaging the score of each cell with the scores of the adjacent cells (Figure 1E).

### Outputs

SURFMAP provides the user with a 2D map in a PNG or PDF format along with a text file containing the mapped values of the map. The text file can be used for numerical comparisons of 2D maps calculated (i) with different features of the same protein thereby enabling the investigation of feature correlations or (ii) with the same feature mapped on different protein homologs.

### Usage

SURFMAP is a very easy-to-use program that only needs as input the pdb file(s) of the protein(s) that the user wants to study. One should notice that if the user aims to compare surface properties of different protein homologs, their 3D structures must be pre-aligned in order to produce comparable 2D maps that are in the same referential. SURFMAP enables the calculation of already encoded features such as electrostatic potential, Kyte-Doolittle or Wimley-White hydrophobicity, stickiness, CV, or can project any feature stored in the temperature factor column of the pdb file (Figure 2).

**Figure 2:**
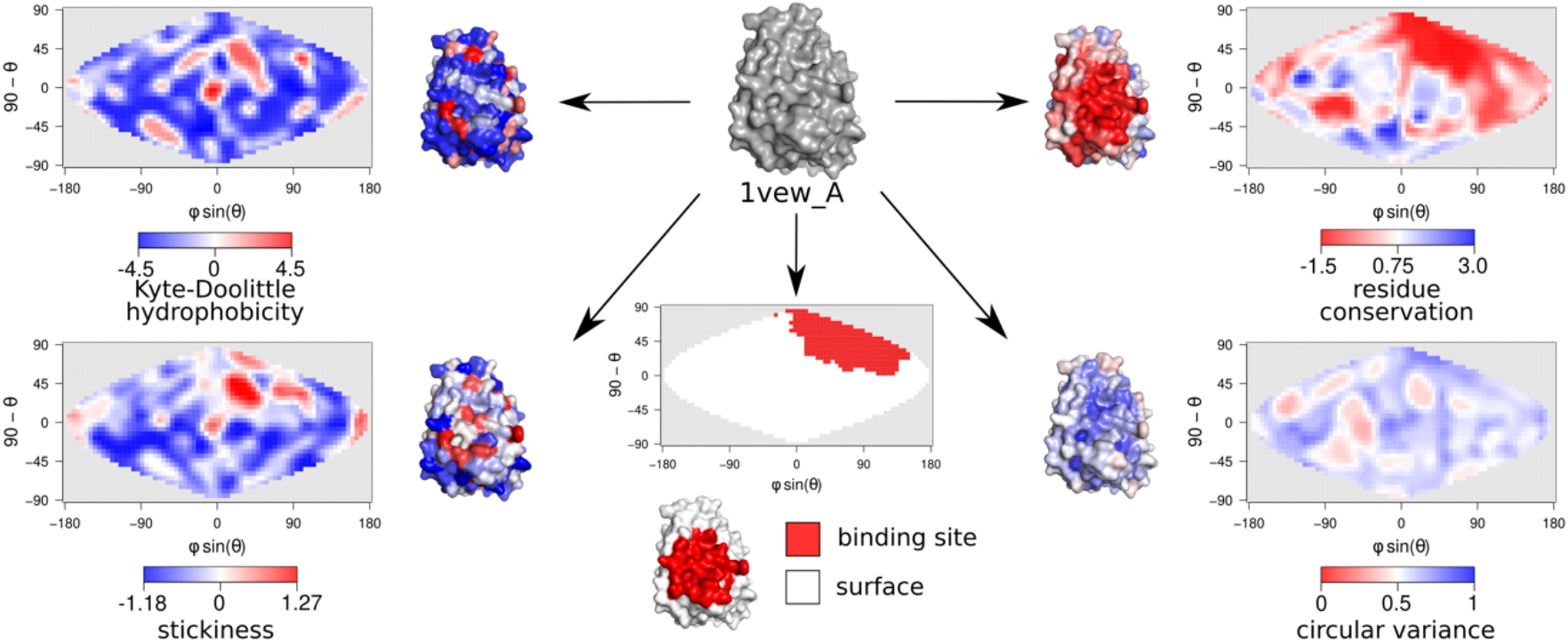
2D maps of different surface features calculated for chain A of the manganese SOD of *E. coli*. The 3D structure of the chain A of 1vew (PDB code: 1vew) is represented in grey (center) along with the corresponding 3D structures and 2D maps colored according to the Kyte-Doolittle hydrophobicity, residue conservation calculated with ConSurf^7^ with conservation scores stored in the temperature column of the input PDB file, stickiness^25^ and circular variance^26^ scales. Dimerization site is mapped with the information stored in the temperature column of the input PDB file (bottom middle map).

SURFMAP is fast and takes on average about 3 seconds to generate a 2D map of a 200 residues globular protein. Electrostatic maps take a longer time (up to 30 seconds for a 200 residues globular protein), the limiting step being the computation of the electrostatic potential with APBS^27^.

## EXAMPLE OF APPLICATIONS ON SUPEROXIDE DISMUTASES

We illustrate the application of SURFMAP with the superoxidase dismutases (SODs), a family of enzymes that catalyze the dismutation of the superoxide radical to hydrogen peroxide and dioxygen. Several types of SODs have been reported according to the metal ions required at the active site. Here we focus on iron (Fe) or manganese (Mn) SODs^30–37^. Some of the Fe and Mn-SODs form homodimers while others form homotetramers, independently of the required metal species. The experimental 3D structures of several SODs have been characterized so far and many studies have attempted to classify them in order to understand the molecular features explaining their Fe and Mn ion specificity, as well as their oligomeric states^31–40^. All these studies rely on the careful examination of sequence alignments or manual inspections of 3D structures.

Here, we computed the 2D maps of several surface descriptors of the chain A of 1vew (1vew_A) which is a subunit of a dimeric manganese SOD. Figure 2 shows the 2D projection of the Kyte-Doolittle hydrophobicity, stickiness, surface relief (i.e., circular variance), and sequence conservation calculated with Consurf^7^, along with the 2D projection of the interaction site participating in the dimer (Figure 2). The 2D maps reveal a heterogeneous distribution of all descriptors with specific patterns distributed over the surface of the protein. The circular variance map is characterized by several red spots that correspond to low circular variance values and indicate protuberant regions. The stickiness reflects the interaction propensity of protein surface regions and is thereby expected to be related to hydrophobicity since protein binding sites are enriched in hydrophobic residues.

Nevertheless, if the stickiness and hydrophobicity maps are overall similar, they display specific patterns and provide complementary information. Indeed, the hydrophobicity map indicates a hydrophobic region (red spot - bottom left) which is not predicted as sticky, but rather displays an intermediate interaction propensity (white spot - bottom left). On the opposite, the stickiness map reveals a sticky spot (red spot on the top right of the map) which does not correspond to a hydrophobic region. Overall, the stickiness map displays a vast non-sticky region (bottom of the map and left part) along with four sticky spots that colocalize on the top right of the map. The latter precisely correspond to the dimer binding site (red spot on the binding site 2D projection map) which is also characterized by a flat and highly conserved region (white region and dark blue spots on the circular variance and sequence conservation maps respectively).

Figure 3 shows how the user can use SURFMAP to compare the surface properties of different homologous proteins. As an example, we compared several surface properties for manganese SODs that either form homodimers (Figure 3 - top) or homotetramers (Figure 3 - bottom). Therefore, we calculated their stickiness, circular variance, and residue conservation 2D maps along with their interaction site 2D projection maps. To produce comparable maps, all proteins were aligned beforehand in the same referential with TMalign^1^. Monomers that participate in dimeric assemblies possess only one interaction site that also exists in the monomers forming tetramers (i.e. common interaction site in red). The tetramers are characterized by two additional binding sites (i.e. tetramer specific binding sites, in yellow and blue respectively), each one involving one of the three other chains of the assembly. Interestingly, if the proteins that dimerize display circular variance maps similar to those of proteins involved in tetramers, their stickiness and sequence conservation maps are clearly different. Indeed, the stickiness maps of the proteins forming dimers are characterized by red spots on the top right of the maps while the proteins forming tetramers display two additional red spots (black stars). The latter, again, correspond to the two additional binding sites involved in the tetramers (blue and yellow binding sites), supporting the stickiness descriptor as a good proxy for probing the interaction propensity of a protein surface. On the other hand, the analysis of the sequence conservation maps reveals for all monomers, a large red spot which locates at the common binding site (red binding site), showing that its amino acids are conserved in both dimers and tetramers. On the other hand, the surface region that corresponds to the blue binding site specific to the tetramers is conserved to a lesser extent in monomers forming tetramers, whereas it is not the case for the corresponding non-binding region in the proteins forming dimers. This reflects that this noninteracting region in proteins forming dimers is not subject to selection. Interestingly, the yellow binding site is not conserved in any of the proteins, including those able to form tetramers. To which extent this binding site is important for tetramerization deserves further investigation.

**Figure 3:**
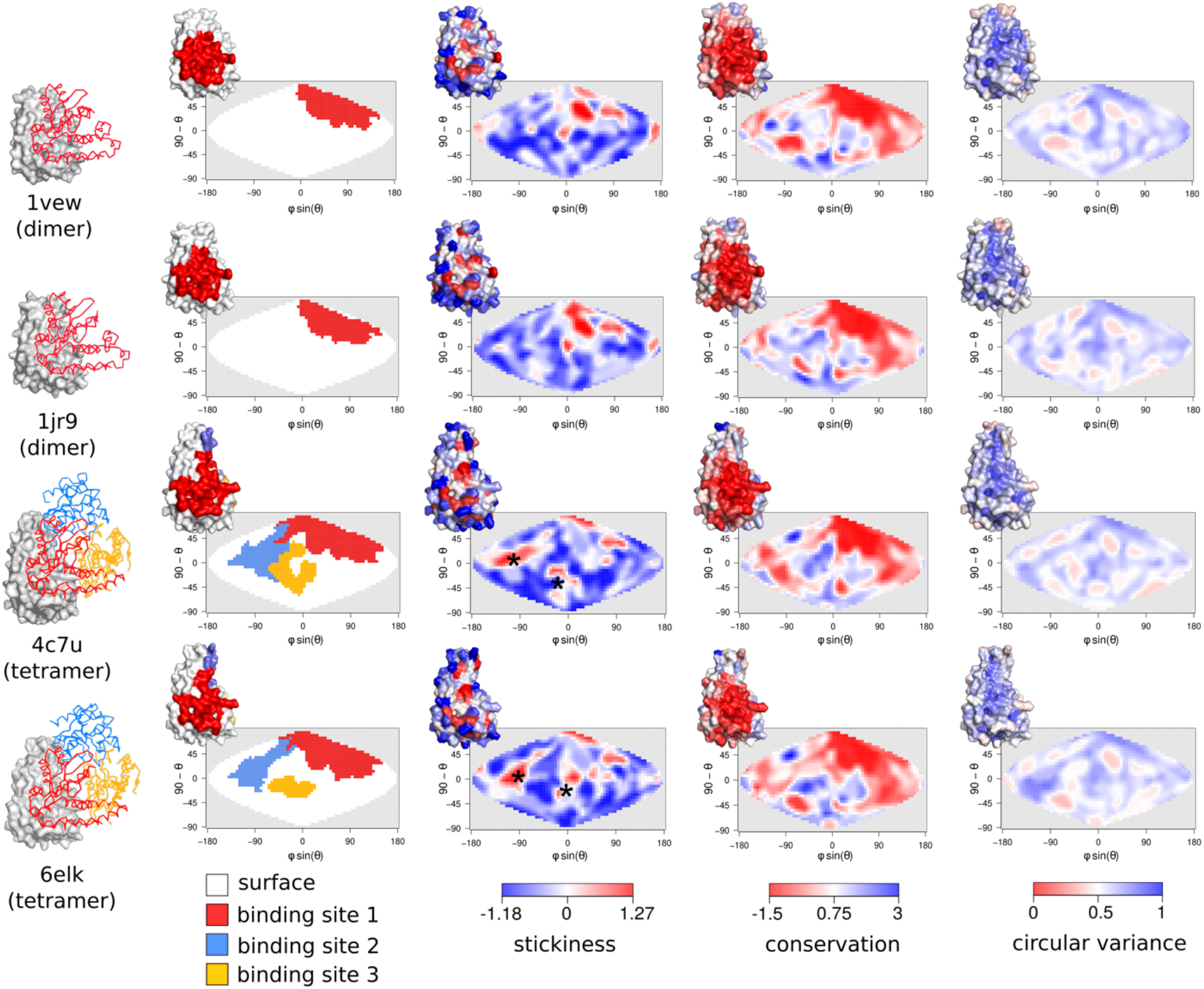
SOD forming homodimers display 2D maps different from those calculated for SODs forming homotetramers. 2D maps calculated for four monomeric chains of manganese SODs that either form dimers (2 first rows - PDB codes: 1vew and 1jr9) or homotetramers (2 bottom rows - PDB codes: 4c7u and 6elk). The 3D structure of each SOD monomer is represented (shown with surface mode of pymol^15^) along with the 3D structures of its corresponding interacting chains (shown with ribbon mode) (one interacting chain colored in red for the two dimeric SODs, and three interacting chains colored in red, blue and yellow respectively for the two tetrameric SODs). For each monomer, are presented their corresponding 2D maps representing the projection of binding site(s) (each binding site is colored according to the involved chain), along with the stickiness, residue conservation calculated with ConSurf, and circular variance distributions over the monomer surface. Each map is also associated with the corresponding surface feature mapped on the 3D structure of the monomer. Black stars indicate sticky spots specific to the two monomers forming tetramers.

In the last section, we show an example of a quantitative comparison of 2D maps calculated for 23 SODs that form either dimers or tetramers (see PDB codes in Figure 4). Therefore, we computed the Manhattan distance between each pair of SOD 2D stickiness maps. For two maps A and B, we summed all Manhattan distances calculated between each pair of corresponding pixels a and b of coordinates (n,m) as follows:

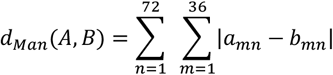

**Figure 4:**
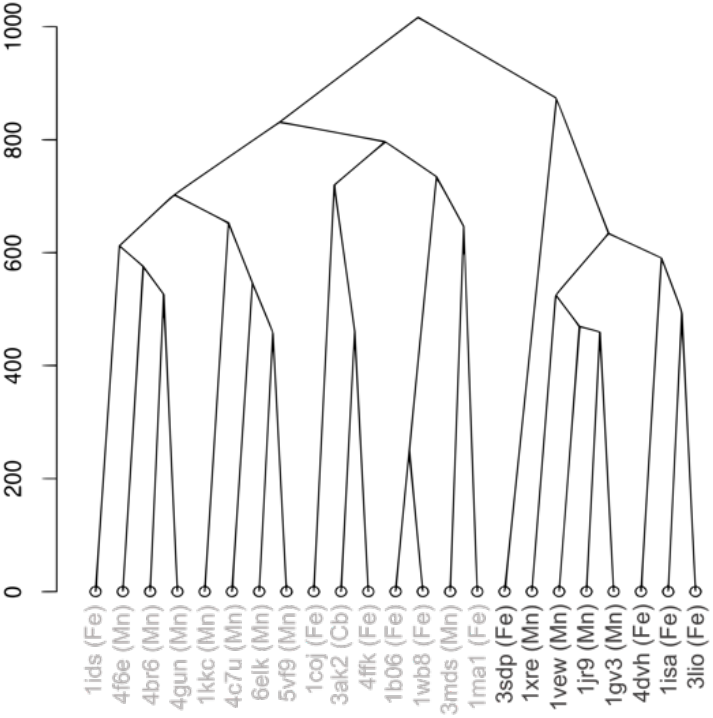
Distance tree based on stickiness 2D map distances. The distance tree is obtained from a hierarchical clustering algorithm (hclust method in R^41^, method = “complete”) based on the stickiness 2D map distance matrix calculated for 23 SODs. PDB codes are indicated for each leaf along with the metal ion preference of each SOD. Monomers forming dimers are indicated in black while those forming tetramers are colored in grey.

Figure 4 shows the distance tree resulting from a Hierarchical clustering based on the map pairwise distances. Interestingly, the distance tree shows two clusters that precisely correspond to monomers forming dimers (black leaves) and tetramers respectively (grey leaves). In addition, in each oligomerization state cluster (dimers or tetramers), the SODs tend to cluster according to their metal binding preference. Interestingly, the distinction between monomers forming dimers or tetramers is less clear in the distance tree calculated from the Kyte-Doolittle hydrophobicity maps, strengthening the stickiness descriptor as a good proxy for comparing different oligomeric states between protein homologs (Figure S1).

## CONCLUSION

We presented SURFMAP, a program that enables the 2D projection of predefined protein surface descriptors or any surface feature stored in the temperature factor of a PDB file of interest. SURFMAP is a free standalone and easy-to-use software that runs on Windows, Linux, and macOS. It provides an easy and visual framework to analyze protein surface features, thereby enabling the user to (i) compare the distribution over the protein surface of different features or (ii) study the evolution of a specific feature(s) across homologous proteins. In addition, SURFMAP provides the map in a text file format allowing the fast and straightforward quantitative comparison of surface features of homologous proteins.

## Supporting Information

Distance tree based on KD hydrophobicity 2D map distances (Figure S1) (PDF).

## Data and Software Availability

SURFMAP can be downloaded freely at https://github.com/i2bc/SURFMAP. SURFMAP can be installed manually or through a docker container. The docker container works on Linux, macOS, and Windows, and is the recommended way to use SURFMAP. The manual installation works only on Linux distributions. SURFMAP requires MSMS^28^, and APBS^27^ for electrostatics calculation. Both are provided in the docker container, but if the user chose the manual installation, the latter must be downloaded and installed separately.

## AUTHOR INFORMATION

### Notes

The authors declare no competing financial interests.

## ACKNOWLEDGMENT

HS and SG works were supported by a French government fellowship. We thank Michel Sanner for authorizing us to integrate MSMS in a docker container. We thank Nathan Baker for authorizing us to integrate APBS in a docker container.

