## Supplementary material for "SURFMAP: a software for mapping in two dimensions protein surface features": Suplemental Material

**Content: 1 supplemental Figure**

**Supplemental Figure S1:** Distance tree based on KD hydrophobicity 2D map distances.

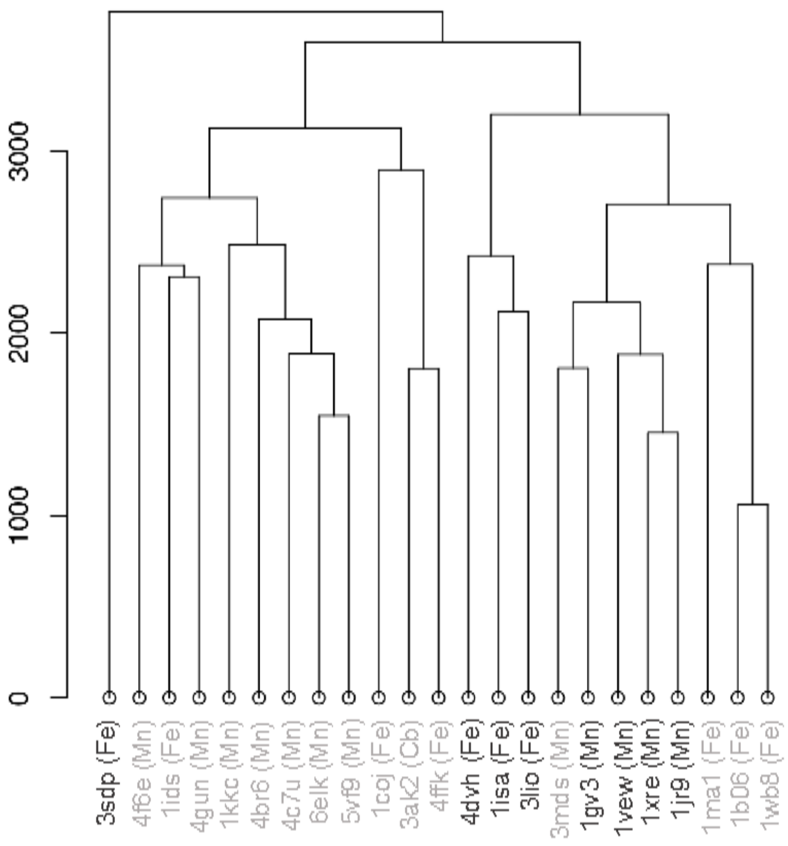


**Figure S1: Distance tree based on KD hydrophobicity 2D map distances.** The distance tree is obtained from a hierarchical clustering algorithm (hclust method in R[1], method = "complete") based on the KD hydrophobicity 2D map distance matrix calculated for 23 SODs. PDB codes are indicated for each leaf along with the metal ion preference of each SOD. Monomers forming dimers are indicated in black while those forming tetramers are colored in grey.
